# IGHV allele similarity clustering improves genotype inference from adaptive immune receptor repertoire sequencing data

**DOI:** 10.1101/2022.12.26.521922

**Authors:** Ayelet Peres, William D. Lees, Oscar L. Rodriguez, Noah Y. Lee, Pazit Polak, Ronen Hope, Meirav Kedmi, Andrew M. Collins, Mats Ohlin, Steven H. Kleinstein, Corey T Watson, Gur Yaari

## Abstract

In adaptive immune receptor repertoire analysis, determining the germline variable (V) allele associated with each T- and B-cell receptor sequence is a crucial step. This process is highly impacted by allele annotations. Aligning sequences, assigning them to specific germline alleles, and inferring individual genotypes are challenging when the repertoire is highly mutated, or sequence reads do not cover the whole V region.

Here, we propose an alternative naming scheme for the V alleles as well as a novel method to infer individual genotypes. We demonstrate the strength of the two by comparing their outcomes to other genotype inference methods and validated the genotype approach with independent genomic long read data.

The naming scheme is compatible with current annotation tools and pipelines. Analysis results can be converted from the proposed naming scheme to the nomenclature determined by the International Union of Immunological Societies (IUIS). Both the naming scheme and the genotype procedure are implemented in a freely available R package (PIgLET). To allow researchers to explore further the approach on real data and to adapt it for their future uses, we also created an interactive website (https://yaarilab.github.io/IGHV_reference_book).

## Introduction

The adaptive immune system’s diversity is key in fighting the array of countless pathogens our bodies encounter. Part of this diversity comes from the immunoglobulin (Ig)-encoding genomic loci, resulting from the stochastic recombination process they undergo. The IG loci are challenging to study because of their repetitive nature and structural variants [19, 46, 31]. In adaptive immune receptor repertoire sequencing (AIRR-seq)-driven studies, a crucial step for downstream analyses is germline annotation, which infers the germline subgroup, gene, and allele for each variable (V), diversity (D), and joining (J) sequence. Studies in the field of adaptive immunity are as diverse as the system itself, yet they need a common language to be able to integrate the data and studies’ conclusions. Understanding of the architecture of the human Ig loci has developed over multiple decades. A widely used taxonomy for human IG genes, which provides a common language for V, D, and J germline subgroups, genes, and alleles [15, 19], was codified by the ImMunoGeneTics Information System (IMGT) [9]. This nomenclature, sometimes referred to as the IMGT nomenclature, is referred to here as the International Union of Immunological Societies (IUIS) nomenclature, for the gene names are allocated according to a process governed by the IUIS. With technological advances in the field, the number of known alleles and genes has increased dramatically [22, 20, 25]. Figure. 1A illustrates the IG heavy chain V (IGHV) locus on chromosome 14, based on the GRCh38 [36] assembly, demonstrating the complexity of the region. A number of genes are duplicated, leading to the presence of genes at different locations (for example IGHV2-70/IGHV2-70D, IGHV3-23/IGHV3-23D) that share common alleles [46]. Additionally, as previously shown for short read IGHV sequences [18], exploring the similarity between all full length functional alleles within the germline set shows that in some cases alleles from different genes are clustered together (Fig. 1B). Closely observing a case of shared alleles between duplicated genes demonstrates the complexity of correctly assigning the germline allele in AIRR-seq data (Fig. 1B, lower panel).

**Figure 1:**
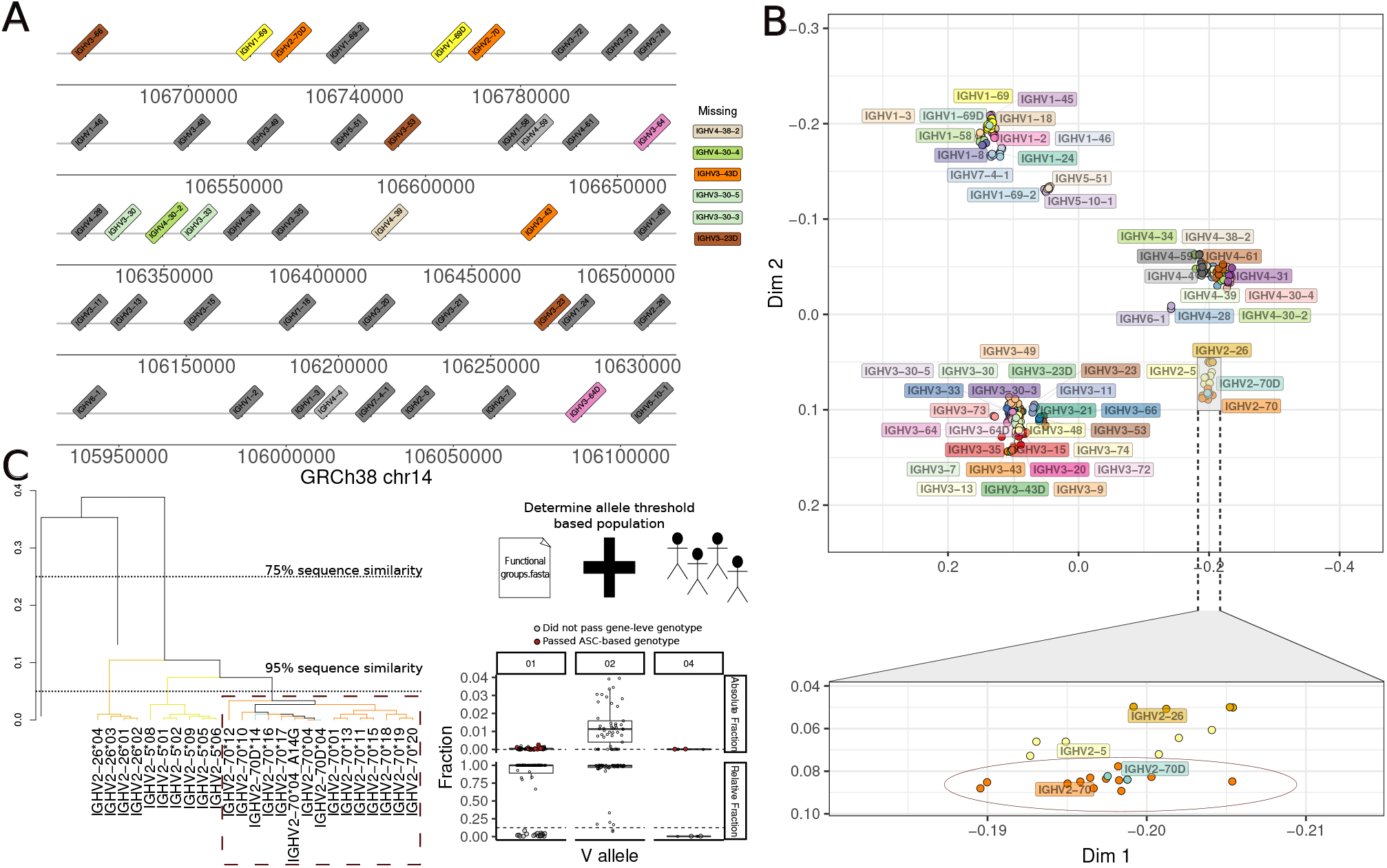
Sequence similarity in the IGH locus. (A) The IGHV locus on human chromosome 14 (x axis, GRCh38 coordinates). The colored genes (non dark-gray) are those with 95%-100% germline sequence similarity. Genes that share similarity, but do not have a genomic location in the GRCh38 assembly, are shown to the right of the plot. Dark gray genes are stand alone genes, and other colors indicate genes with similar alleles. (B) Multidimensional scaling of the IGHV IMGT germline set pairs distance matrix; plot shows the first two dimensions. Each dot is a functional allele colored by gene. The bottom panel shows a zoom into the IGHV2 subgroup, demonstrating the proximity between alleles of the duplicated genes (IGHV2-70 and IGHV2-70D). (C) The left panel shows the hierarchical clustering of the IGHV2 subgroup. The color of the branches was determined by the gene color in panel B. The right panel is a schematic representation of the new ASC-based approaches for genotyping. The top and bottom panels show the genotype inference for a given ASC/gene. The columns are the alleles of the ASC/gene, and the rows are the different genotyping method. Each dot is an individual’s allele call frequency, calculated appropriate to the genotyping method (Top panel relative to the total repertoire size, and bottom panel to the ASC/gene size). The dotted lines represent the genotype thresholds: the allele specific threshold (1e-04) in the top panel, and the gene-based threshold (0.125) in the lower panel. The gray and red points represent individual allele calls that did not enter the genotype based on the gene-based method, but did based on the ASC-method (respectively).

Germline annotation is typically performed by an aligner tool, which determines the germline allele by comparison to sequences listed in a ’germline set’. For V genes, the accuracy of this assignment is strongly influenced by the sequencing read length [25, 49]. Reads that cover the entire V sequence (typically 290-320nt in length) permit the greatest accuracy, but shorter reads are often employed, and many studies focus only on sequencing the complementarity-determining region 3 (CDR3) with short flanking sequences, thus dramatically reducing the number of alleles that can be categorically resolved, particularly as there is reduced diversity at the 3’ end of the V gene germline sequences [28, 26, 25]. Even when full-length reads of the V sequence are available, sequence alignment against the germline set will not provide a single categorical germline allele for every sequence, both because of duplicated sequences in the germline set itself, and because even a small number of mismatches from the germline can cause a V sequence to become equidistant from *>* 1 sequence in the set. As a result, the aligner tool will frequently emit ’multiple assignments’ a list of germline alleles that statistically indistinguishable to the V sequence present in the read.

This complexity of assignments impacts clonal inference [49] among other things. Clones are a measure of diversity and selection within B cell receptor (BCR) repertoires [48]. Each clone stems from an ancestral naive B cell expressing an unmutated BCR. In AIRR-seq repertoires, it is common to identify a BCR clone as a group of sequences that share the same V and J germline assignments, and CDR3 length [10], as well as having similarity in the CDR3 sequence. To achieve correct clonal inference, annotating the AIRR-seq data correctly is therefore crucial, and mis-assignments and multiple assignments can result in difficulty to infer clones, hindering any clonal-based downstream analysis. For example, in Fig. 1C, 14 alleles between IGHV2-70 and IGHV2-70D have a nucleotide sequence similarity greater than 95%.

To address this problem, we propose the use of a naming scheme system for analysis that is based on the hierarchical clustering of alleles, according to nucleotide sequence similarity. In our system, gene families are defined as sequences with 75% similarity [19], and ’allele similarity clusters’ (ASCs) as groups of sequences that share 95% similarity (Fig. 1C left panel). Essentially, assignments are made to ASCs rather than genes for the purposes of downstream analysis. Sequence similarity is based here on germline sequences that are matched to read length, hence the number of clusters, and overall precision of the annotation, reflects the precision of the underlying germline set. The 95% threshold represents the best relation between clustering similar alleles and avoiding “over” splitting known groups (Genes). At the end of analysis, where it is necessary to refer to specific allele assignments, the alleles names can be converted to the familiar IUIS nomenclature: we do not propose a replacement nomenclature, but rather a method of data representation that is more tractable for analysis.

Because of the high sequence identity between many alleles, aligner tools typically infer a biologically implausible number of alleles in an individual’s repertoire [6]. Tools inferring ‘personal genotypes’ assess support for each allele [7, 30, 34]. This can be helpful in downstream analyses such as disease susceptibility inference [2, 21]. We propose a genotyping approach based on consideration of the absolute expression of each allele, using ASC annotation to ensure that each allele sequence is only considered once (Fig. 1C right panel). We show that this provides improved results compared to a commonly used existing tool and is in good correspondence with genotypes derived from genomic sequencing.

## Results

### Allele naming system based on germline hierarchical structure

Using hierarchical clustering with complete linkage, we defined a two-level naming scheme for the set of functional germline alleles (downloaded from IMGT July 2022): allele families (AFs) and ASCs. For the family level, we followed the logic and threshold (75% nucleotide similarity) from IMGT [17]. Since we applied this methodology to the contemporary set of functional alleles, the resulting families mildly deviate from the present IUIS family definitions. In particular, the IGHV3 subgroup is split into two families with our approach (Fig. 2A, the orange dashed circle defines the 75% threshold). Using the same hierarchical tree, we clustered the sequences based on 95% nucleotide similarity (Fig. 2A, blue dashed line). This resulted in 46 clusters, which we define as ASCs, some of which span several genes (Fig. 2B). In addition, alleles of some genes are split between different clusters. Adapting the two-level naming scheme results in an annotated germline reference set that reduces ambiguities in several analysis steps as shown below.

**Figure 2:**
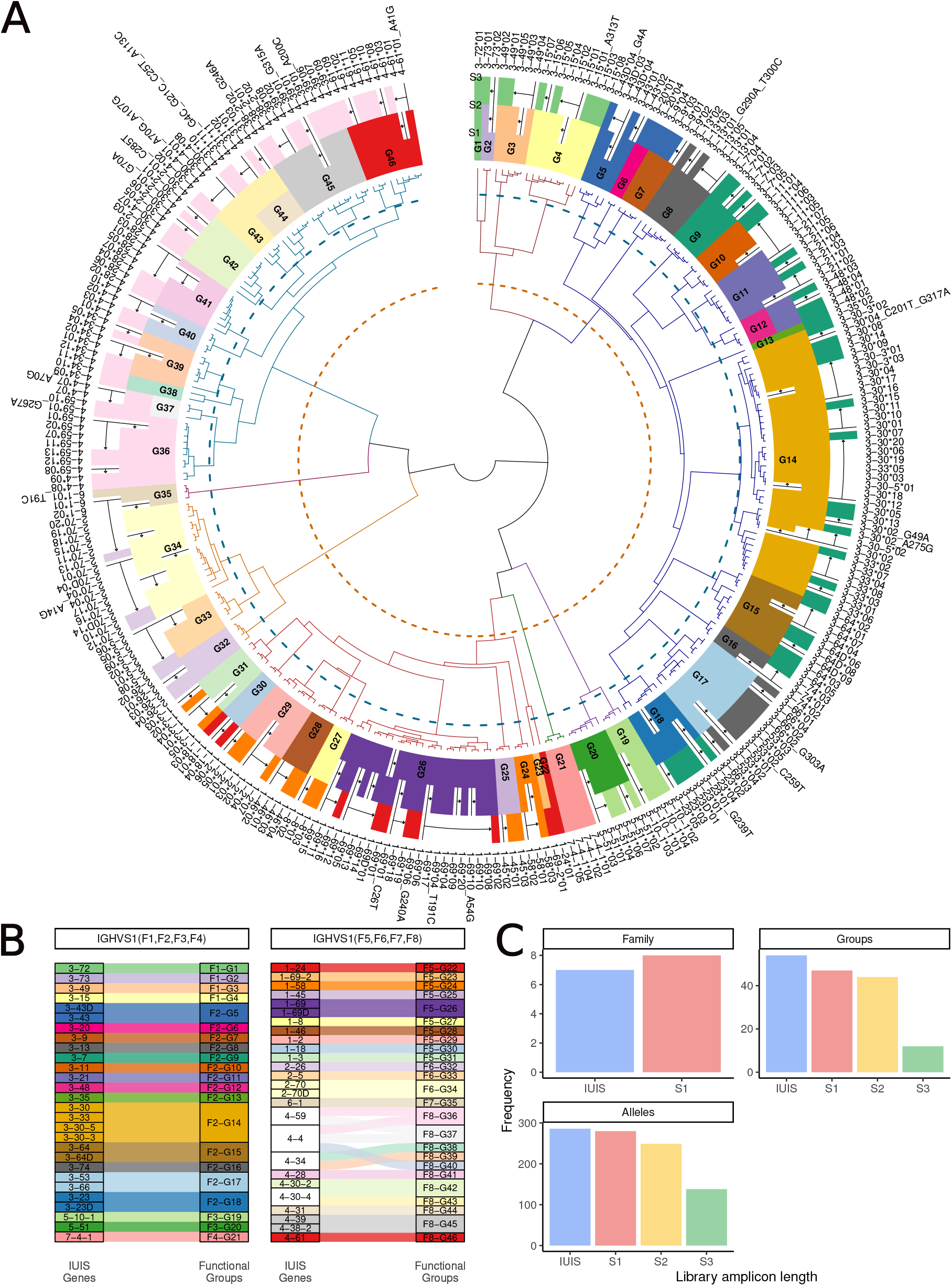
Allele Similarity Clusters. (A) Hierarchical clustering of the functional IGH germline set. The inner layer shows a dendrogram of the clustering, the dotted circles indicate the sequence similarity of 75% (orange) and 95% (blue). The dendrogram branches are colored by the 75% sequence similarity. The first colored circle shows the clusters and alleles for the library amplicon length of S1, the second circle for the length of S2, and the third for S3. The white color indicates alleles that cannot be distinguished in the library’s germline set. (B) An alluvial plot showing the connection between the allele clusters and the IUIS genes. The colors represent the allele clusters. White represents IUIS genes whose alleles are clustered into more than a single allele cluster. (C) The frequency of the subgroups/families, genes/clusters, and alleles for each amplicon length. The x-axis is the amplicon length and the y-axis is the count of the unique subgroups/families, genes/clusters, or alleles.

Many AIRR-seq experimental protocols result in sequences that do not cover the full V gene. The two most common partial sequencing libraries are BIOMED-2 [39], with primers in the framework 1 (FW1) and framework 2 (FW2) regions, and ImmunoSeq [23], which offers only the CDR3 and a small fragment of the V and J region. Partial V sequencing exacerbates the computational challenges mentioned above caused by similar alleles originating from distinct genes. Our proposed naming scheme can be generalized in a straightforward way to these situations.

To adapt the above naming scheme to partial V sequences, we computationally trimmed the 5’ region of the germline set’s V sequences according to the sequence lengths obtained using the BIOMED-2 and ImmunoSeq protocols. For simplicity, we defined the sequencing protocols by the library amplicon length, and named the full-length amplicons “S1”, the partial V sequences corresponding to the BIOMED-2 style “S2”, and the minimal V coverage of ImmunoSeq “S3” (Fig. 2A).

Depending on the amplicon length used, we obtained a different number of ASCs. As expected, the 5’ V trimming resulted in higher similarity between the alleles. Compared to the 54 genes in the IUIS database, after clustering we observed 46 ASCs in S1, 43 in S2, and 11 in S3 (Fig. 2C).

### Allele similarity clusters based threshold for genotyping are robust to nomenclature and more accurate than gene-based inference

Many computational genotyping tools consider the relative frequency of a candidate allele during inference and filtering steps [7, 5, 37, 3]. An inference is made or accepted if the number of assignments to an allele exceeds a threshold percentage of the total assignments to all alleles of the corresponding gene. A need for allele-specific filtering processes has been suggested in the past [24]. Here, we propose and implement a method based on the explicit comparison of each allele’s frequency in the repertoire under study with that observed in the population. We do so by using the ASC naming scheme so that each identical nucleotide sequences belonging to different alleles are collapsed and represented in the germline reference set only once (Fig. 1C right panel). This addresses issues that can confound frequency observations in current methods: variable expression levels between alleles of the same gene, multiple assignments, duplicated genes, and short reads. In our implementation, we initially set a default allele threshold of 1*e*^*−*^4 for each of the alleles in the ASC germline set. We then manually adjusted the threshold, such that each allele has an allele-specific threshold (Sup. Table 1). This was determined from observations of the allele’s usage across all available naive B cell samples present in VDJbase [26]. In particular, thresholds were adjusted in the following cases: A. haplotype inference suggested that the default threshold led to a non-sensible biological scenario. B. The allele usage in a given individual was very far from the usage distribution of across the whole sampled population. Overall, 129 of 280 thresholds were adjusted. For further refining the threshold from specific populations of interest, we developed an interactive web server ([https://yaarilab.github.io/IGHV_reference_book/]). The server presents the frequencies of the alleles and the chosen thresholds. Further, the server allows the end user to explore different choices for the allele threshold and inspect the implications of these modifications by comparing matching haplotype data for available individuals. We found such modification to be helpful in maximizing discrimination, particularly in cases where some alleles of the gene are found to have lower expression levels than others [14, 8, 28, 24].

After reviewing and, where necessary, adjusting the allele-specific thresholds for all the IGHV alleles observed in two naive cell datasets (VDJbase projects P1 [8] and P11 (unpublished study), 142 repertoires in total), we compared the resulting inferred genotypes with the ones inferred by TIgGER, a gene-based inference tool (Fig. 3A). In the ASC-based genotype approach, alleles enter the genotype if their usage is higher than the allele-specific thresholds. In TIgGER genotype inference on the other hand, the alleles enter the genotype based on the relative usage normalized by all sequences mapped to this gene. A common step in this kind of analysis includes an undocumented allele inference. In the ASC-based genotype inference, each inferred undocumented allele is given the allele-specific threshold of its most similar allele. Overall, there were 5695 allele calls that were included in either or both genotypes. Results were concordant for inference of highly used alleles, with 5471 allele calls that fully matched between the methods (95%, green squares). However, there were 4 allele calls that only entered the genotype with the gene-based method (pink squares), and 220 alleles that were called only by the ASC-method (black squares).

**Figure 3:**
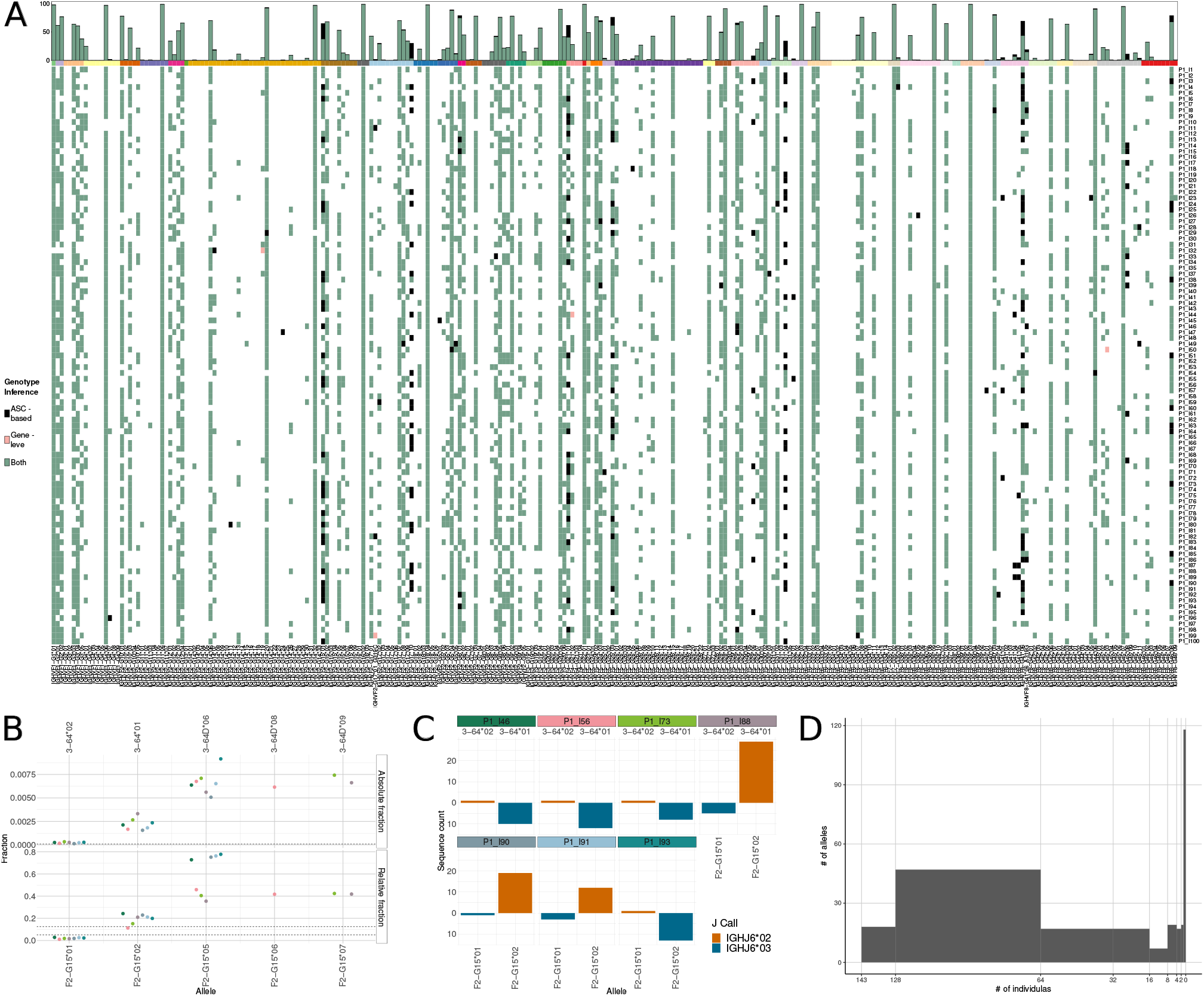
Genotype comparison. (A) A heatmap comparing the genotypes inferred from the gene-based and the ASC-based method. The bottom panel is the heatmap comparison, where each row is a genotype inference of an individual from the P1 dataset and each column is a different allele. Black and Pink colors represent alleles that only entered the genotype either in the ASC-based method or in gene-based method with a 12.5% threshold, respectively. Green represents alleles that entered the genotype in both methods, and white represents alleles that did not pass in both methods. The top panel is the summation of the heatmap events. The y-axis is the count of the individuals for which a given allele entered the genotype. The x-axis is the different alleles. (B) The relative and absolute frequency of the ASC IGHVF2-G15. Each dot is an individual for which the allele 01 entered the genotype with the ASC-based method, but did not in the gene-based method. The colors represent the different individuals. Each column is a different allele from the cluster. The top row is the absolute frequency and the bottom is for the relative frequency. (C) Haplotype based on IGHJ6 for the individuals from (B). Each facet is a different individual, and the facet color matches the dots from (B). In each facet, the top row and orange color is the frequency for the IGHJ6*02 chromosome and the bottom and blue color for the IGHJ6*03 chromosome. The x-axis is the different alleles for the cluster, and the y-axis is the frequency of the sequence count. (D) The distribution of the allele abundance in the population. The x-axis is the number of individuals attributed to each allele, and the y-axis is the number of alleles.

The potential false positives in the gene-based genotyping were seen in cases where all observed alleles of a particular gene were expressed at low levels, according to population data. An example is IGHV7-4-1 (part of ASC IGHVS1F4-G21). In all individual genotype inferences, there was not a single situation of heterozygosity for this cluster, as in most individuals there was one dominant allele. In the single occasion where heterozygosity was declared, both alleles *01 and *02 entered the genotype. However, the inference of allele *02 is likely to be incorrect. In this particular sample (VDJbase: P1 I44 S1), the poorly expressed allele was *01, with 4 sequences, while the highly expressed allele *02 only had a single sequence. This deviates from what is seen in the population. The three alleles attributed to this cluster vary in usage, with allele *02 being the most expressed allele [24] with a median usage of 1.19*e*^*−*02^. This is 33 times more than the second expressed allele (*01). Hence, the situation where allele *01 dominates over allele *02 is unlikely (p value of 4 *×* 10^*−*6^ according to a binomial test), and the identification of a read associated to allele *02 might be the result of a sequencing error. This indicates clear deviations between the approaches that may lead to different specificities in lowly expressed clusters.

Potential false negatives in the gene-based genotyping are seen in cases where one allele is expressed at a lower rate than other alleles of the gene, according to population data. An example is IGHV3-64*02 (corresponding to ASC IGHVS1F2-G15*01). This allele entered the genotype using the ASC-based method, but not using the gene-based method. The IGHVS1F2-G15 cluster combines alleles from two IUIS genes, IGHV3-64 and IGHV3-64D, that merge under the 95% threshold. The alleles of this cluster vary in usage, i.e., alleles *05, *06, and *07 are more frequently used than 02 and 01 (Fig. 3B). Allele IGHVS1F2G15*01 is expressed at a considerably lower rate than the other alleles, with a median of 2.1*e*^*−*04^ absolute usage: roughly 12 times lower than the second most lowly expressed allele, IGHVS1F2-G15*02 (aka IGHV3-64*01). Even so, allele IGHVS1F2-G15*01 (aka IGHV364*02) was above the ASC-based threshold (1*e*^*−*04^) and entered the genotype, while being far below the gene-based relative fraction threshold of 12.5% or 5%, with a median of 1.86% (3B). To validate the inference of allele IGHVS1F2-G15*01, we looked at the haplotype of alleles IGHVS1F2-G15*01 and IGHVS1F2-G15*02, since they come from the same chromosomal location (IGHV3-64). We haplotyped seven individuals who ostensibly included allele *01 with the ASC-method but not in the gene-based method, using heterozygosity at IGHJ6 as the anchor (Fig. 3C). In all seven individuals, alleles *01 and *02 are found on opposite chromosomes, strongly supporting the presence of allele IGHVS1F2-G15*01. This example demonstrates the sensitivity of the ASC-based approach to lowly expressed allele inferences, which may provide important insights in future studies.

Fig. 3D summarizes the distribution of allele prevalence in cohorts P1 and P11 (Fig. S1). Seven out of the 280 alleles present in the ASC germline reference set appear in all 142 individuals, while 41% of the alleles, 116 out of 280, do not enter any of the genotypes. This could imply that reference sets should be population-specific [43, 27], or that the current reference set includes a large fraction of unexpressed or non-existent alleles [35].

### Allele usage reporting

Subgroups, genes, and sometimes alleles are commonly used as AIRR-seq features, for example in reporting over-expression of specific genes/families in the context of specific diseases. These features are highly sensitive to the nomenclature and the genotypes of the individuals in the cohort. Here, we compare the reporting of allele-level usage versus gene or cluster level. Fig. 4 shows that reporting of usage is highly influenced by the genotypes of the individuals. For example, in IGHVS1F2-G5 the mean absolute usage of individuals who carry alleles *04 and *05 is significantly higher than those who carry *03 and *04. If we were to report the overall ASC instead of the usage of the carried allele combinations, the mean usage would have been closer to the lowly expressed combination, masking the differences. Moreover, if IUIS genes were used to report the usage of these alleles, it would have been split between the V3-43 and V3-43D columns, as the genes share a common allele that would have resulted in a multiple assignment in the alignment process. Consequently, when studying allele usage in human cohorts, we recommend that usage is reported at the ASC level, to avoid unnecessary ambiguities.

**Figure 4:**
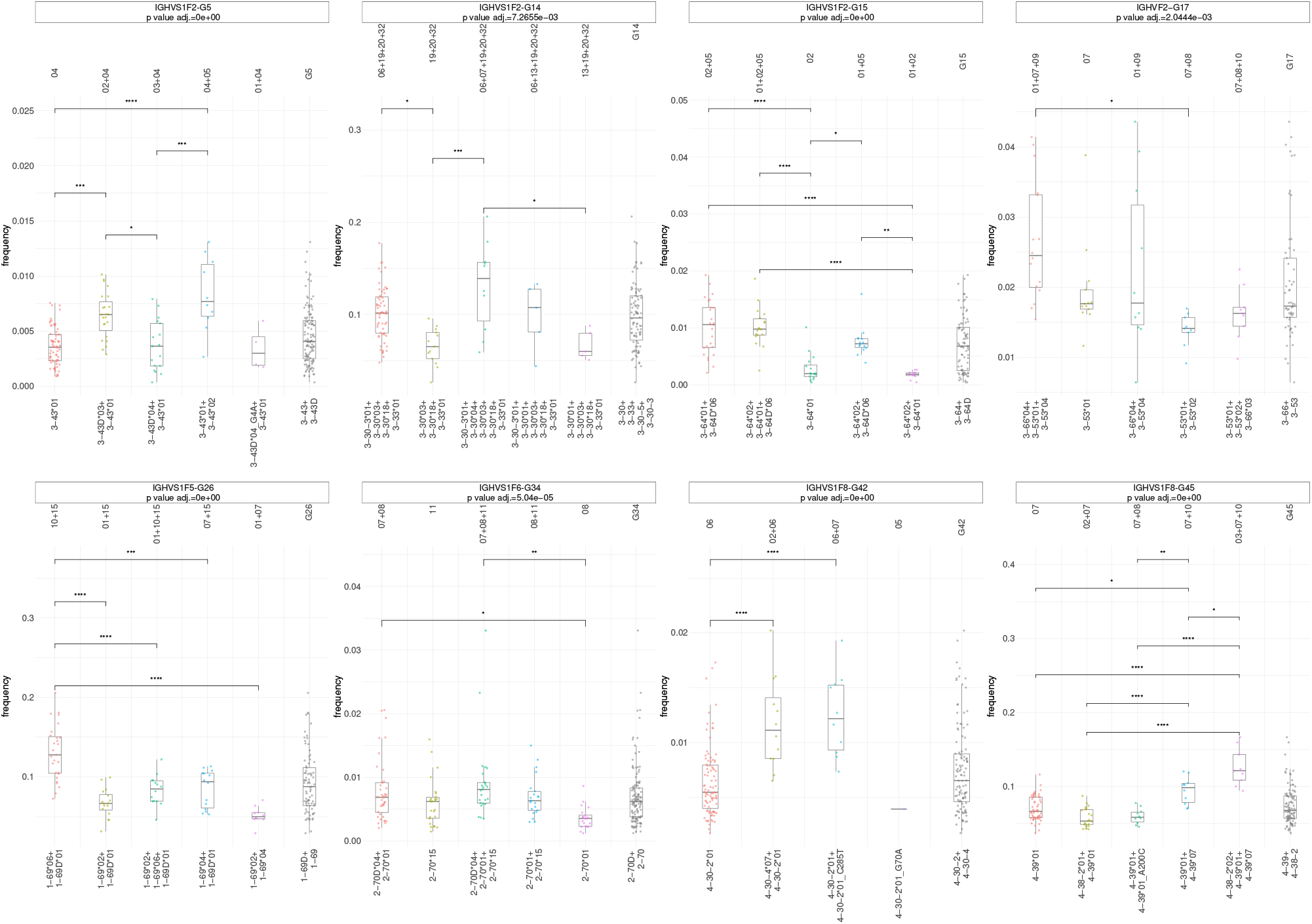
Gene usage-based genotype. The absolute usage frequency (calls out of the total repertoire size) of the top five allele combinations for the given clusters. The x-axis columns are the cluster’s allele combination after genotype inference, ordered by the number of individuals carrying the combinations. The y-axis is the absolute frequency of the cluster within the total repertoire. Each point is an individual’s absolute frequency. The colors represent the order of the combinations, where the combination which is present in most individuals is colored in red and so on. The gray color, last column in every x-axis facet, indicates the absolute frequency in terms of the whole cluster. For each cluster, an ANOVA test was calculated and the adjusted p value is presented in the cluster’s plot title. A Tukey’s HSD multiple comparison test was calculated with the adjusted p value, comparing between allele clusters indicated on the connecting line; only the statistically significant combination were drawn. ns: *p >* 0.05, *: *p <*= 0.05, **: *p <*= 0.01, ***: *p <*= 0.001, ****: *p <*= 0.0001

**Figure 5:**
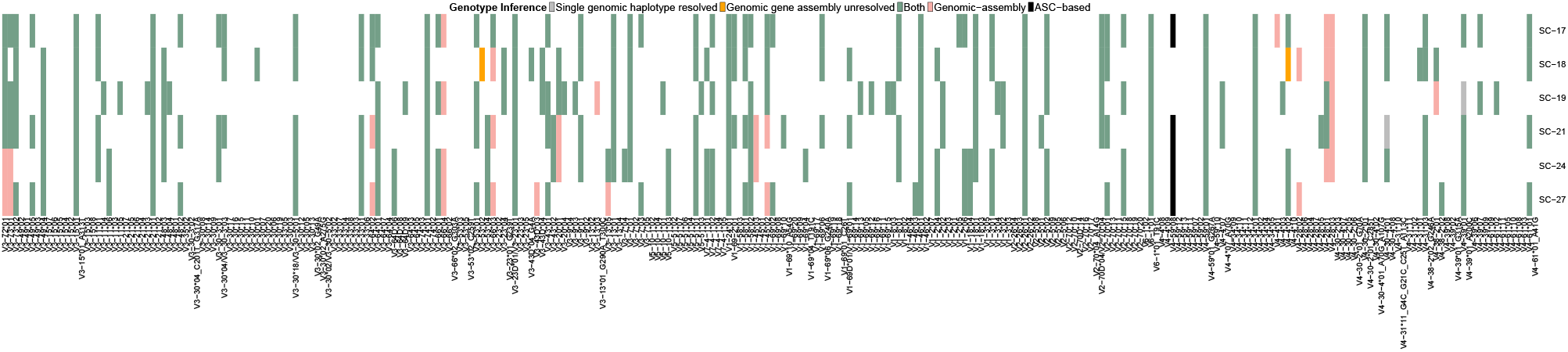
Genomic inference compared to AIRR-seq genotype inference. A heatmap of a comparison between the ASC-based genotype method and genomic validation. Each row is an individual, and each column is a different allele. Black and Pink colors represent alleles that are only present in the ASC-based method or in genomic validation, respectively. Green represents alleles that are present in both methods, and white represents alleles that are not seen in either. Orange represents alleles that for them there are no gene evidence in the genomic validation, and gray represents alleles that only one haplotype was resolved in the genomic validation.

### Genomic validation of the ASC-based genotype

We validated our ASC-based genotyping method using a paired dataset drawn from six subjects, comprising AIRR-seq repertoire sequencing of IgM naive enriched cells, and a haplotyped assembly of the genomic IGHV locus derived from long-read sequencing (REF). Across the six subjects (5), a total of 304 ASC allele calls were made from the AIRR-seq repertoires (counting the identification of a single allele in a single individual as an allele call).

In several subjects, the genomic assembly was incomplete, either not covering certain genes at all, or not resolving to haplotypes in certain regions. In total, 2 of the 304 calls were in genes without coverage (orange squares), and 2 in locations with unresolved haplotypes (gray squares). This left 301 allele calls from the repertoires that could be verified in the assemblies. Of those calls, 296 (*>* 97 percent) were concordant between the ASC and genomic results (green squares). In the four discordant cases, the assemblies for the particular gene were resolved, but only a single allele was observed (black squares). This contrasted with the ASC inferred genotype, in which donors were characterized as heterozygous, carrying two alleles. All cases were of the allele IGHV4-59*08.

34 allele calls were found only in the genomic samples (pink squares) and not in the ASC genotypes, implying that these alleles are poorly or not at all expressed. Such examples have been described in the literature [24, 16]

In summary, out of the 304 allele calls that were made across six individual genotypes using the ASC-based method, we found potential contradictions from the genomic data only in four cases. These cases most likely indicate technical issues with the genomic assembly due to reduced coverage, rather than in the ASC-based genotype inference method.

### Generalizability to other germline sets

One potential limitation of the proposed naming scheme is that the specific alleles in the germline reference set determine the allele families and ASCs, hence the clustering may change when alleles are added or removed from the set. To quantify the impact of an altered germline reference set, we created a reduced germline set consisting only of the alleles that entered the genotype of P1 individuals, as determined by our ASC-based method. This is an example of transferring one germline reference set from one dataset to another without adjusting it. We then applied the clustering algorithm and obtained the new families and clusters (Fig. 6A). Compared to the original set, two cluster pairs were merged, G36/G37 and G43/G44, and G13 and G38 were dropped, as none of their alleles entered the genotype. The overall structure was maintained, despite the reduction from 280 to 163 alleles (Fig. 6B). From this, we conclude that the clustering method is relatively robust to changes in the reference set composition.

**Figure 6:**
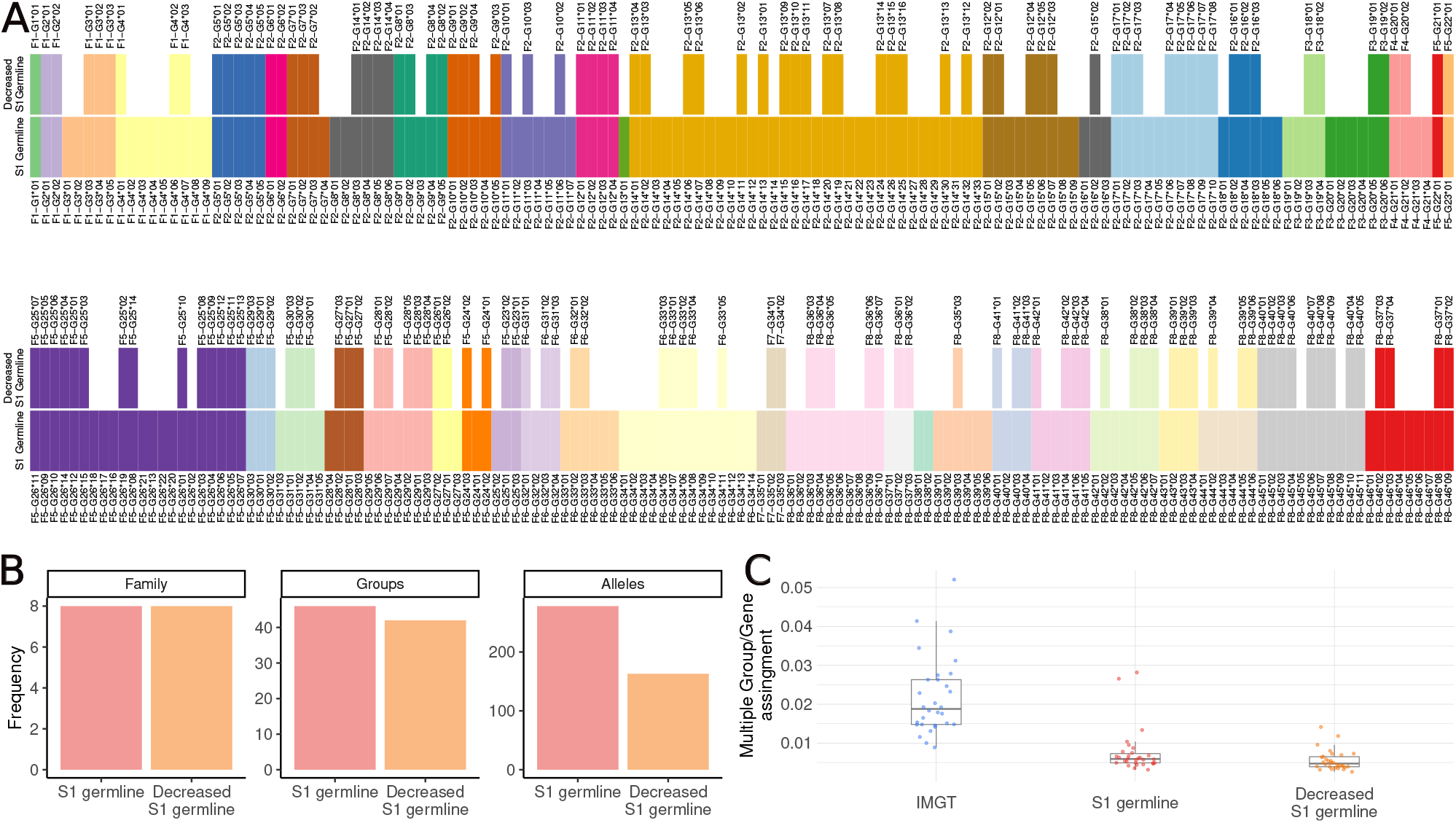
Reduced germline-based genotype. (A) The heatmap shows the clusters based on the full germline used in Figure 2, and the re-clustering after the reduced germline, which includes alleles that entered the genotype using the ASC-based method on the P1 and P11 cohorts. The bottom row shows the clusters for the full germline set, and the top row shows the clusters for the new germline. The colors represent the different clusters. White represents alleles that did not enter the genotype. (B) Summation of the number of families, clusters, and alleles in the S1 germline and the reduced reference. The x-axis is the different reference sets and the y-axis is the count of the events. (C) The frequency of multiple cluster/gene assignments. The x-axis is the different reference sets and the y-axis is the absolute frequency of multiple assignments. Each dot is an individual’s multiple assignment frequency from the non-naive P4 cohort.

To further assess the flexibility and effect of the reference set, we tested the multiple assignments in a non-naive repertoire. Multiple assignments are cases in which the aligner, IgBlast in our evaluations, cannot determine the single matched allele, and outputs multiple options for the most likely germline ancestor allele. This can be caused by sequencing errors, somatic hypermutation, identical germline alleles shared by multiple genes, or a combination. We explored this effect using the P4 dataset from VDJbase, which includes non-naive repertoires from 28 individuals. We aligned the repertoires three times, once with the IUIS gene definitions downloaded from IMGT (the IMGT set), once with an identical set of sequences but using the proposed assignment nomenclature (the S1 set), and once with the reduced germline set described above (the reduced S1 set). We calculated the fraction of sequences that were attributed by the aligner to more than a single gene/ASC. Figure 6C shows an expected but significant reduction of 3-fold in multiple assignments between the IMGT set and the S1 set. The reduced S1 set showed a further reduction in multiple assignments.

We then applied the ASC approach to other AIR-encoding genomic loci. We clustered the sets of functional alleles downloaded from IMGT (July 2022) for human IGKV, IGLV, TRBV, and TRAV. We applied the same thresholds of 75% and 95% for determining the allele families and ASCs (Fig. 7). The IGK locus is unique because of its duplicated pattern. The locus has two V gene blocks with a large gap in between, where the 3’ distal block is essentially an inverted duplication of the 5’ block. Here, as in IGHV, some genes share alleles with identical sequences. As expected, these duplicated alleles are clustered together under the 95% threshold. A split is observed in IGKV1-17, whose alleles are assigned to two ASCs. In the IGL locus, where IUIS defines 10 subgroups, we found 12 families using our approach and thresholds. Four genes were combined into ASCs, and a single gene was split into two ASCs. The loci of TRB and TRA remained relatively constant, except for four TRBV genes, which were merged into two ASCs. We developed an interactive application that applies the ASC naming scheme to V allele reference sets from different loci and organisms, https://yaarilab.github.io/IGHV_reference_book/alleles_groups.html. As the reference set can change over time, we recommend not to use the nomenclature in reporting but only in the downstream analyses. Nevertheless, for backtracking, reproducibility, and interoperability, we maintain an https://doi.org/10.5281/zenodo.7401189 of all ASC runs conducted by our web server. Allowing translation of the allele cluster names into IUIS names and also into the unique names suggested in the supplementary materials (Sup. Table 1).

**Figure 7:**
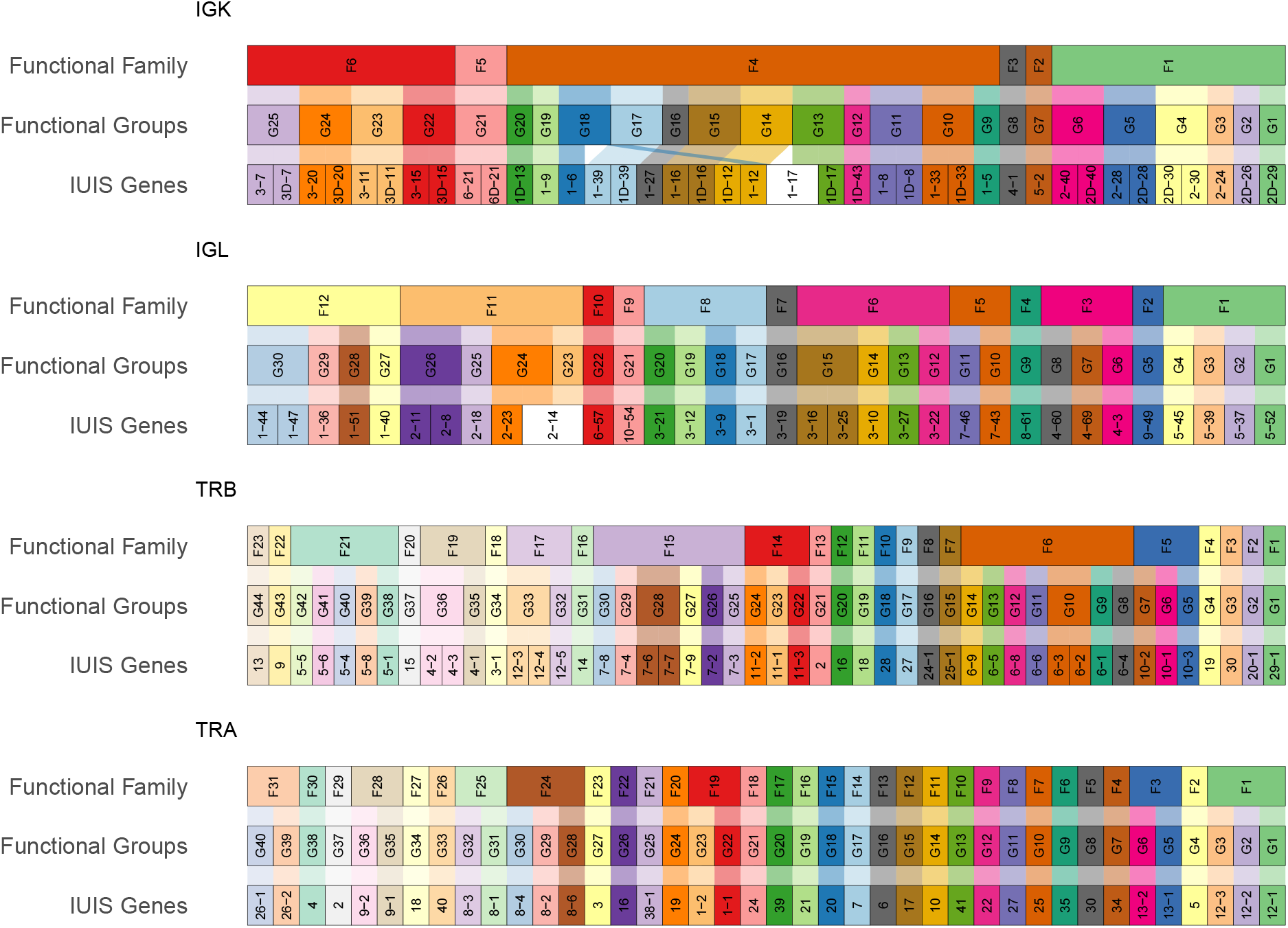
Allele clusters for V genes from IGL/K and TRB/A loci. Each alluvial plot represents the cluster division for a given locus. The first row of each plot shows the division of the families, the second row the ASCs, and the third IUIS gene clustering. The colors represent the allele clusters. White represents IUIS genes that have been re-clustered into more than a single allele cluster.

## Discussion

Several groups have used repertoire sequencing to study IG and TR loci, using inferencing tools. They discovered a plethora of undocumented allele sequences [5, 8, 25, 20, 22, 40, 12, 45, 38]. With careful review, many novel alleles identified in the human loci may be mapped to specific genes, on the basis that their sequence clusters closely with other alleles of a single gene. [13, 24]. In other species, the genes are not well characterized, and macaque and mouse germline sets resulting from these studies are published as discrete sets of allele sequences unmapped to genes. This can pose challenges when trying to genotype and haplotype using the conventional method, which is based on the gene level. In this study, we report on two innovations that can be highly beneficial in such situations. The first is our proposed naming scheme that organizes the alleles within clusters of sequence similarities, which aids downstream analyses. The ASCs can be used for clonal inference, usage reporting, and genotype and haplotype inference. We believe that the ASC naming scheme can be a good starting point until more information on the layout of the species’ genomic loci is discovered. That being said, the proposed scheme is not meant to replace the existing IUIS naming, but rather an accompanied set to allow for a more inclusive analysis. We created an R package (PIgLET, https://bitbucket.org/yaarilab/piglet), and an online application within the ASC website (https://yaarilab.github.io/IGHV_reference_book/alleles_groups.html) that allows the users to infer the ASC based on their own V allele reference set and plot the ASC results. Another potential use case for our proposed naming scheme is clonal inference. Many of the clonal inferences rely on the V segment assignments, which can be influenced by similar genes and alleles. Considering this factor in the clonal inference can be influential on the final results. Therefore, utilizing the ASC approach may lead to better clustering tools.

The second innovation we report is a new and improved approach for inference of a personal genotype and for determination of VDJ gene usage from AIRR-seq data. The approach is based on the absolute frequency of allele usage within a specific population, rather than on relative usage (normalized at the gene level), as other approaches do. We created an interactive website where each page shows the allele usage across the naive IGHV repertoires from P1 and P11 studies of VDJbase. The site allows users to play with the data and explore the ASC-base thresholds. Further, the website includes an interactive interface to create ASCs based on a reference set. Our site will be continuously updated as more naive AIRR-seq and direct genomic sequencing datasets accumulate. Along with the site, the thresholds for allele detection in VDJbase will also be updated. Moreover, as new species are sequenced and published, we will include them as well in the site and in VDJbase. It is worth mentioning a potential issue with all AIRR-seq based genotype approaches: in some rare cases two alleles differ only at the 3’ end of the sequence (in human IGHV, *>* 318), imposing many instances of multiple assignments as the aligners cannot differentiate between the two when the rearrangements are trimmed before. In human IGHV, only two such cases exist (3-66*01 and 3-66*04, 4-28*01 and 4-28*03). These cases should be treated separately, considering all particularities of the sequences, and should be reported with an adequate confidence level.

We have demonstrated the application of ASC-based allele usage information to the analysis of over- or under-expression in specific diseases or conditions. Annotation with ASCs tailored to the sequencing read length employed, followed by ASC-based genotyping, will provide a single orthogonal vector of allele usages that can be compared across repertoires, eliminating the complexity and bias that can arise from the much larger numbers of multiple assignments produced by gene-based approaches. The allele usage vector provides a clear signal, tailored to the precision of the underlying data set, which can be used in graphical analysis or machine learning applications. Important conclusions can be translated back to IUIS nomenclature.

It is known that some alleles of a gene may be expressed at higher levels than other alleles. Gene-based genotyping based on transcriptomic data can overlook relatively lowly expressed alleles, however, our ASC-based method, which takes account of the typical levels of allele expression, will add them to the genotype correctly. Identifying lowly expressed alleles and including them in the genotype can be critical for investigating disease susceptibility [41, 42, 11, 47]. Since genotypes are relatively similar within populations [27], variations in susceptibility to diseases are plausibly caused by such small differences [2].

We validated the ASC-based approach by comparing AIRR-seq genotypes with a genotype based on direct long read genomic sequencing [31]. Even though some repertoires in these genomically sequenced cohorts had relatively low AIRR-seq depths, the comparison showed a strong concordance between the direct sequencing and the proposed inference method.

Our results show reduced variability in genotypes among individuals, as compared with the current IGH reference available in IMGT. This raises an interesting debate of whether all alleles in the existing reference set truly exist. This point was previously reviewed in [44], in which the authors discovered that several alleles were erroneous. We believe that this matter should be further discussed and reviewed to curate an optimal reference set for AIRRseq analyses. Exploring naive repertoires is far from complete, as most studies focus on the same ethnic populations. As demonstrated by Rodriguez et al. [32] different ethnicity influence the IGH composition (i.e genes, deletions, etc.). With more repertoire data curated with different ethnic background, the allele specific threshold might have to be tailored toward the ethnic population. We envisage that with the rising interest in AIRR-seq, future studies will provide more diversity, which will contribute to the efforts to enhance both the ASC website and VDJbase, and to optimization of the inferences and tools.

## Methods

### Data

Naive and non-naive BCR repertoire heavy chain data were used, of individuals from three VDJbase [26] projects, P1, P11 (naive), and P4 (non-naive). Library preparation and processing for projects P1 and P11 were performed as described in [8]. The processing for project P4 repertoires is described in [4]. The most recent IMGT IGHV reference set was downloaded in July 2022. For this study, we downloaded the V, D, and J allele reference set from IMGT on July 2022, the reference set included the functionality annotation for each allele. Within the V reference set, alleles which were non-functional were discarded. This lead to discarding subgroup IGHV8, as none of the alleles in this subgroup are functional. Hence, the V reference set includes only alleles from subgroups IGHV1-7.

### ASCs

To create the ASCs, we used the most recent available IGHV reference set from IMGT, with addition of undocumented allele sequences inferred from both P1 and P11 cohorts. The combined set was then filtered to include only functional alleles that start from the first position of the V sequence region (as defined by IMGT numbering scheme). We then discarded short sequences in the 5’ end, those that do not start in the first nucleotide, and short sequences in the 3’ end: the upper limit was chosen based on the quantile that contains the largest number of sequences with the longest coverage. The position selected was 318, shown to be a reliable position for inferring undocumented alleles [20].

To cluster the alleles, we calculated the Levenshtein distance between all allele pairs after aligned to the IMGT numbering scheme. For calculating the distance, we trimmed the sequences to the 3’ upper limit position of 318. We then used hierarchical clustering with complete linkage. The final tree was cut by two similarity thresholds of 75% and 95% to obtain the allele families and ASCs. As a result of the clustering, the alleles were renamed to represent the new allele families and ASCs (Supp. Table S1, [29]). For example, in the allele IGHVS1F2-G15*02, the family is represented by F2, the ASC by G15, and the allele by 02. The S1 is an indicator of the library amplicon length of a given reference set. A key table that links between the IUIS naming scheme and the ASC naming scheme can be found in the supplementary (Sup. Table 1). The reference set with the new naming scheme was then used for downstream processing.

For the downstream analysis, we used the full length of the V germline sequence. Meaning, without the 3’ trim that was used for the clustering. As the alleles 01 and 04 of IGHV3-66, and alleles 01 and 03 of IGHV4-28, only differentiate in position 319 it is collapsed in the cluster analysis. Hence, in the reference set we have decided to add both alleles for both genes, however in the new name scheme allele IGHV3-66*04 is marked as a novel allele.

### Allele similarity clusters based genotype method

The ASC-based genotype utilizes a population derived thresholds to determine the presence of a given allele within an individual’s genotype. We first had to set a default threshold for the absolute allele usage fraction before tailoring it to each allele, the chosen value was 10^*−*4^. We then observed the absolute usage of each allele within its ASC to determine the final allele-specific threshold. A list of all thresholds can be found here (Sup. Table S1).

### AIRR-seq processing

For inferring the personal genotype either by the ASC-based or the gene-based method, the repertoires were first aligned with IgBlast (V1.17) using the customized germline set. Then, for each clone a representative with the least number of mutations was chosen, undocumented alleles were inferred using TIgGER [7], and in case new alleles were found, the repertoire was realigned. Then, if the dataset came from naive B cells, the sequences were filtered for no mutations within the V region up to position 316, accounting for possible sequencing errors at the end of the V region [8]. For repertoires coming from full V region length amplicons, the repertoires were filtered to omit 5’ trimmed sequences. Sequences were also filtered for sufficient 3’ coverage of the V region (at least 312 nucleotide long). For the ASC-based inference, the allele’s absolute usage was calculated, and in cases where a given sequence had more than a single assignment, the counts were divided among all clusters. For each allele within the repertoire, the absolute usage was compared to the specific threshold. Alleles that passed the threshold were then added to the final individual’s genotype. For the gene-based genotype inference, the TIgGER ’inferGenotype’ [7] function was used with the chosen threshold, either 12.5% or 5%.

### Using genome long-read assemblies to validate alleles

Long-read assemblies from 6 samples generated using SequelIIe HiFi reads and Igenotyper were downloaded from Rodriguez et al. [33]. The assemblies were aligned to a custom immunoglobulin heavy chain (IGH) genome reference containing previously discovered IGHV genes using BLASR [46, 1]. From the alignments, gene sequences (5’ UTR, leader-1, leader-2 and exons) were extracted. The matched repertoire sequences were processed as described in the method section (AIRR-seq processing), reducing the small fraction of nonnaive cells in the following way: after the initial alignment and the inference of novel alleles, we inferred clones using the new allele clusters, and then for each clone we chose the least mutated sequence as a representative. Because the sequences were pre-sorted to IgM, we inferred the genotype only for unmutated sequences (after inferring novel alleles).

## Acknowledgements

We thank all members of the iARC sub-committee of the AIRR community for productive discussions. This study was partially supported by grants from the ISF (2940/21), the United States–Israel Binational Science Foundation (2017253), VATAT grant, and the European Union’s Horizon 2020 research and innovation program (825821 to GY). The contents of this document are the sole responsibility of the iReceptor Plus Consortium and can under no circumstances be regarded as reflecting the position of the European Union.

