## Supplementary material for "IGHV allele similarity clustering improves genotype inference from adaptive immune receptor repertoire sequencing data"

[1]Faculty of Engineering, Bar Ilan University, 5290002 Ramat Gan, Israel
[2]Bar Ilan Institute of Nanotechnology and Advanced Materials, Bar Ilan University, 5290002 Ramat Gan, Israel
[3]Institute of Structural and Molecular Biology, Birkbeck College, University of London, London, United Kingdom
[4]Department of Biochemistry and Molecular Genetics, University of Louisville School of Medicine, Louisville, KY, United States
[5]Yale School of Medicine, New Haven, Connecticut, USA.
[6]Division of Hematology and Bone Marrow Transplantation, Chaim Sheba Medical Center, Tel-Hashomer, Israel
[7]Sackler School of Medicine, Tel-Aviv University, Tel-Aviv, Israel
[8]The Mina & Everard Goodman Faculty of Life Sciences, Bar-Ilan University, Israel
[9]School of Biotechnology and Biomedical Sciences, University of New South Wales, Sydney, Australia
[10]Department of Immunotechnology, Lund University, Lund, Sweden

December 26, 2022

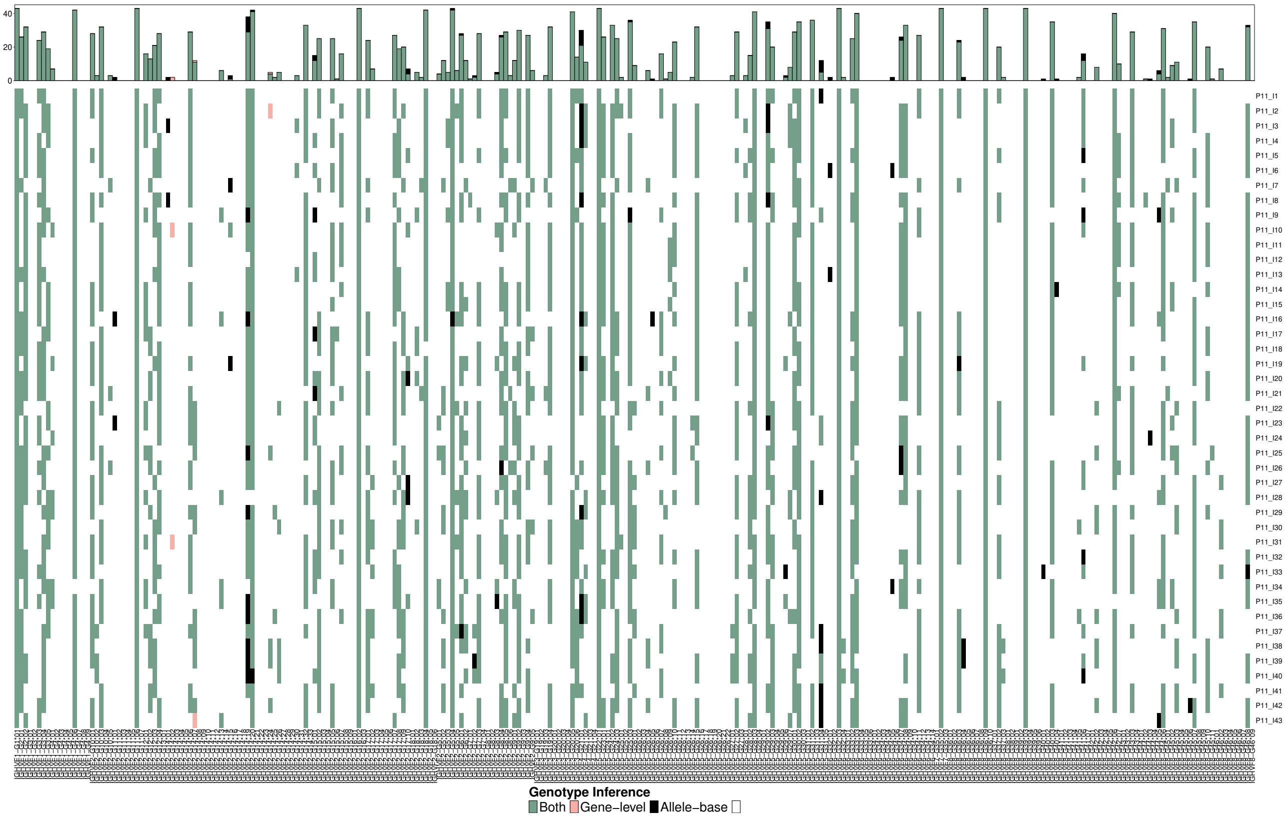


Figure S1: **P11 individuals genotype comparison.** A heatmap comparing the genotypes inferred from the gene-level and the allele-based method. The bottom panel is the heatmap comparison, where each row is a genotype inference of an individual from the P11 dataset and each column is a different allele. Black and Pink colors represent alleles that only entered the genotype either in the allele-based method or in gene-based method with a 12.5% threshold, respectively. Green represents alleles that entered the genotype in both methods, and white represents alleles that did not pass in both methods. The top panel is the summation of the heatmap events. The y-axis is the count of the individuals for which a given allele entered the genotype. The x-axis is the different alleles.

| ASC Allele | IUIS Allele | Allele threshold |
|---|---|---|
| IGHVF1S1-G1*01 | IGHV3-72*01 | 1e-04 |
| IGHVF1S1-G2*01 | IGHV3-73*01 | 1e-04 |
| IGHVF1S1-G2*02 | IGHV3-73*02 | 1e-04 |
| IGHVF1S1-G3*01 | IGHV3-49*02 | 1e-04 |
| IGHVF1S1-G3*02 | IGHV3-49*01 | 1e-04 |
| IGHVF1S1-G3*03 | IGHV3-49*05 | 1e-04 |
| IGHVF1S1-G3*04 | IGHV3-49*03 | 1e-04 |
| IGHVF1S1-G3*05 | IGHV3-49*04 | 1e-04 |
| IGHVF1S1-G4*01 | IGHV3-15*07 | 0.001 |
| IGHVF1S1-G4*02 | IGHV3-15*06 | 1e-04 |
| IGHVF1S1-G4*03 | IGHV3-15*05 | 0.001 |
| IGHVF1S1-G4*04 | IGHV3-15*04 | 0.001 |
| IGHVF1S1-G4*05 | IGHV3-15*02 | 1e-04 |
| IGHVF1S1-G4*06 | IGHV3-15*01 | 0.001 |

| | | |
|---|---|---|
| IGHVF1S1-G4*07 | IGHV3-15*01_A313T | 0.001 |
| IGHVF1S1-G4*08 | IGHV3-15*03 | 1e-04 |
| IGHVF1S1-G4*09 | IGHV3-15*08 | 1e-04 |
| IGHVF2S1-G10*01 | IGHV3-11*01 | 0.001 |
| IGHVF2S1-G10*02 | IGHV3-11*04 | 0.001 |
| IGHVF2S1-G10*03 | IGHV3-11*06 | 1e-04 |
| IGHVF2S1-G10*04 | IGHV3-11*03 | 1e-04 |
| IGHVF2S1-G10*05 | IGHV3-11*05 | 0.001 |
| IGHVF2S1-G11*01 | IGHV3-21*07 | 1e-04 |
| IGHVF2S1-G11*02 | IGHV3-21*05 | 0.001 |
| IGHVF2S1-G11*03 | IGHV3-21*06 | 0.001 |
| IGHVF2S1-G11*04 | IGHV3-21*04 | 0.001 |
| IGHVF2S1-G11*05 | IGHV3-21*03 | 0.001 |
| IGHVF2S1-G11*06 | IGHV3-21*01 | 0.001 |
| IGHVF2S1-G11*07 | IGHV3-21*02 | 0.001 |
| IGHVF2S1-G12*01 | IGHV3-48*03 | 0.001 |
| IGHVF2S1-G12*02 | IGHV3-48*04 | 0.001 |
| IGHVF2S1-G12*03 | IGHV3-48*01 | 0.001 |
| IGHVF2S1-G12*04 | IGHV3-48*02 | 0.001 |
| IGHVF2S1-G13*01 | IGHV3-35*02 | 1e-04 |
| IGHVF2S1-G14*01 | IGHV3-30-3*02 | 1e-04 |
| IGHVF2S1-G14*02 | IGHV3-30*04_C201T_G317A | 0.001 |
| IGHVF2S1-G14*03 | IGHV3-30*08 | 1e-04 |
| IGHVF2S1-G14*04 | IGHV3-30*14 | 1e-04 |
| IGHVF2S1-G14*05 | IGHV3-30*09 | 1e-04 |
| IGHVF2S1-G14*06 | IGHV3-30-3*01 | 1e-04 |
| IGHVF2S1-G14*07 | IGHV3-30*04/IGHV3-30-3*03 | 0.001 |
| IGHVF2S1-G14*08 | IGHV3-30*17 | 1e-04 |
| IGHVF2S1-G14*09 | IGHV3-30*16 | 0.001 |
| IGHVF2S1-G14*10 | IGHV3-30*15 | 1e-04 |
| IGHVF2S1-G14*11 | IGHV3-30*11 | 1e-04 |
| IGHVF2S1-G14*12 | IGHV3-30*10 | 0.001 |
| IGHVF2S1-G14*13 | IGHV3-30*01 | 1e-04 |
| IGHVF2S1-G14*14 | IGHV3-30*07 | 0.001 |
| IGHVF2S1-G14*15 | IGHV3-30*20 | 1e-04 |
| IGHVF2S1-G14*16 | IGHV3-30*06 | 1e-04 |
| IGHVF2S1-G14*17 | IGHV3-30*19 | 1e-04 |
| IGHVF2S1-G14*18 | IGHV3-33*05 | 0.001 |
| IGHVF2S1-G14*19 | IGHV3-30*03 | 0.01 |
| IGHVF2S1-G14*20 | IGHV3-30*18/IGHV3-30-5*01 | 1e-04 |
| IGHVF2S1-G14*21 | IGHV3-30*12 | 1e-04 |
| IGHVF2S1-G14*22 | IGHV3-30*05 | 1e-04 |
| IGHVF2S1-G14*23 | IGHV3-30*13 | 1e-04 |
| IGHVF2S1-G14*24 | IGHV3-30*02_G49A | 0.001 |
| IGHVF2S1-G14*25 | IGHV3-30*02_A275G | 0.001 |

| | | |
|---|---|---|
| IGHVF2S1-G14*26 | IGHV3-30*02/IGHV3-30-5*02 | 0.001 |
| IGHVF2S1-G14*27 | IGHV3-33*02 | 1e-04 |
| IGHVF2S1-G14*28 | IGHV3-33*07 | 1e-04 |
| IGHVF2S1-G14*29 | IGHV3-33*04 | 1e-04 |
| IGHVF2S1-G14*30 | IGHV3-33*08 | 0.001 |
| IGHVF2S1-G14*31 | IGHV3-33*03 | 1e-04 |
| IGHVF2S1-G14*32 | IGHV3-33*01 | 1e-04 |
| IGHVF2S1-G14*33 | IGHV3-33*06 | 0.001 |
| IGHVF2S1-G15*01 | IGHV3-64*02 | 1e-04 |
| IGHVF2S1-G15*02 | IGHV3-64*01 | 1e-04 |
| IGHVF2S1-G15*03 | IGHV3-64*07 | 0.001 |
| IGHVF2S1-G15*04 | IGHV3-64*04 | 1e-04 |
| IGHVF2S1-G15*05 | IGHV3-64D*06 | 1e-04 |
| IGHVF2S1-G15*06 | IGHV3-64D*08 | 1e-04 |
| IGHVF2S1-G15*07 | IGHV3-64D*09 | 1e-04 |
| IGHVF2S1-G15*08 | IGHV3-64*03 | 1e-04 |
| IGHVF2S1-G15*09 | IGHV3-64*05 | 1e-04 |
| IGHVF2S1-G16*01 | IGHV3-74*03 | 0.001 |
| IGHVF2S1-G16*02 | IGHV3-74*01 | 1e-04 |
| IGHVF2S1-G16*03 | IGHV3-74*02 | 1e-04 |
| IGHVF2S1-G17*01 | IGHV3-66*01 | 0.001 |
| IGHVF2S1-G17*01_G319C | IGHV3-66*04 | 0.001 |
| IGHVF2S1-G17*02 | IGHV3-66*02 | 1e-05 |
| IGHVF2S1-G17*03 | IGHV3-66*02_G303A | 1e-04 |
| IGHVF2S1-G17*04 | IGHV3-53*03 | 1e-04 |
| IGHVF2S1-G17*05 | IGHV3-53*05 | 1e-04 |
| IGHVF2S1-G17*06 | IGHV3-53*02_C259T | 0.001 |
| IGHVF2S1-G17*07 | IGHV3-53*01 | 0.001 |
| IGHVF2S1-G17*08 | IGHV3-53*02 | 0.001 |
| IGHVF2S1-G17*09 | IGHV3-53*04 | 1e-04 |
| IGHVF2S1-G17*10 | IGHV3-66*03 | 1e-05 |
| IGHVF2S1-G18*01 | IGHV3-23*02 | 1e-04 |
| IGHVF2S1-G18*02 | IGHV3-23*04 | 0.001 |
| IGHVF2S1-G18*03 | IGHV3-23*01_G239T | 0.001 |
| IGHVF2S1-G18*04 | IGHV3-23D*01/IGHV3-23*01 | 0.001 |
| IGHVF2S1-G18*05 | IGHV3-23*03 | 1e-04 |
| IGHVF2S1-G18*06 | IGHV3-23*05 | 1e-04 |
| IGHVF2S1-G5*01 | IGHV3-43D*04_G4A | 1e-04 |
| IGHVF2S1-G5*02 | IGHV3-43D*03 | 1e-04 |
| IGHVF2S1-G5*03 | IGHV3-43D*04 | 1e-04 |
| IGHVF2S1-G5*04 | IGHV3-43*01 | 1e-04 |
| IGHVF2S1-G5*05 | IGHV3-43*02 | 1e-04 |
| IGHVF2S1-G6*01 | IGHV3-20*01 | 1e-04 |
| IGHVF2S1-G6*02 | IGHV3-20*04 | 1e-04 |
| IGHVF2S1-G7*01 | IGHV3-9*04 | 1e-04 |

| | | |
|---|---|---|
| IGHVF2S1-G7*02 | IGHV3-9*03 | 1e-04 |
| IGHVF2S1-G7*03 | IGHV3-9*01 | 0.001 |
| IGHVF2S1-G7*04 | IGHV3-9*02 | 0.001 |
| IGHVF2S1-G8*01 | IGHV3-13*02 | 1e-04 |
| IGHVF2S1-G8*02 | IGHV3-13*03 | 0.001 |
| IGHVF2S1-G8*03 | IGHV3-13*01_G290A_T300C | 1e-04 |
| IGHVF2S1-G8*04 | IGHV3-13*05 | 1e-04 |
| IGHVF2S1-G8*05 | IGHV3-13*01 | 1e-04 |
| IGHVF2S1-G8*06 | IGHV3-13*04 | 1e-04 |
| IGHVF2S1-G9*01 | IGHV3-7*04 | 0.001 |
| IGHVF2S1-G9*02 | IGHV3-7*01 | 0.001 |
| IGHVF2S1-G9*03 | IGHV3-7*02 | 0.001 |
| IGHVF2S1-G9*04 | IGHV3-7*03 | 0.001 |
| IGHVF2S1-G9*05 | IGHV3-7*05 | 1e-04 |
| IGHVF3S1-G19*01 | IGHV5-10-1*02 | 1e-04 |
| IGHVF3S1-G19*02 | IGHV5-10-1*04 | 1e-04 |
| IGHVF3S1-G19*03 | IGHV5-10-1*01 | 0.001 |
| IGHVF3S1-G19*04 | IGHV5-10-1*03 | 0.001 |
| IGHVF3S1-G20*01 | IGHV5-51*02 | 5e-04 |
| IGHVF3S1-G20*02 | IGHV5-51*07 | 5e-04 |
| IGHVF3S1-G20*03 | IGHV5-51*06 | 5e-04 |
| IGHVF3S1-G20*04 | IGHV5-51*04 | 5e-04 |
| IGHVF3S1-G20*05 | IGHV5-51*01 | 5e-04 |
| IGHVF3S1-G20*06 | IGHV5-51*03 | 5e-04 |
| IGHVF4S1-G21*01 | IGHV7-4-1*01 | 1e-05 |
| IGHVF4S1-G21*02 | IGHV7-4-1*02 | 1e-04 |
| IGHVF4S1-G21*03 | IGHV7-4-1*04 | 1e-04 |
| IGHVF4S1-G21*04 | IGHV7-4-1*05 | 1e-04 |
| IGHVF5S1-G22*01 | IGHV1-24*01 | 1e-04 |
| IGHVF5S1-G23*01 | IGHV1-69-2*01 | 1e-04 |
| IGHVF5S1-G24*01 | IGHV1-58*03 | 0.001 |
| IGHVF5S1-G24*02 | IGHV1-58*01 | 1e-04 |
| IGHVF5S1-G24*03 | IGHV1-58*02 | 1e-04 |
| IGHVF5S1-G25*01 | IGHV1-45*03 | 1e-05 |
| IGHVF5S1-G25*02 | IGHV1-45*01 | 1e-05 |
| IGHVF5S1-G25*03 | IGHV1-45*02 | 1e-05 |
| IGHVF5S1-G26*01 | IGHV1-69*02 | 0.001 |
| IGHVF5S1-G26*02 | IGHV1-69*08 | 0.001 |
| IGHVF5S1-G26*03 | IGHV1-69*10 | 0.001 |
| IGHVF5S1-G26*04 | IGHV1-69*10_A54G | 0.001 |
| IGHVF5S1-G26*05 | IGHV1-69*20 | 1e-04 |
| IGHVF5S1-G26*06 | IGHV1-69*09 | 0.001 |
| IGHVF5S1-G26*07 | IGHV1-69*04 | 0.001 |
| IGHVF5S1-G26*08 | IGHV1-69*04_T191C | 0.001 |
| IGHVF5S1-G26*09 | IGHV1-69*17 | 0.001 |

| | | |
|---|---|---|
| IGHVF5S1-G26*10 | IGHV1-69*06 | 0.001 |
| IGHVF5S1-G26*11 | IGHV1-69*06_G240A | 0.001 |
| IGHVF5S1-G26*12 | IGHV1-69*19 | 1e-04 |
| IGHVF5S1-G26*13 | IGHV1-69*18 | 0.001 |
| IGHVF5S1-G26*14 | IGHV1-69*01_C26T | 0.001 |
| IGHVF5S1-G26*15 | IGHV1-69D*01/IGHV1-69*01 | 0.001 |
| IGHVF5S1-G26*16 | IGHV1-69*14 | 0.001 |
| IGHVF5S1-G26*17 | IGHV1-69*13 | 0.001 |
| IGHVF5S1-G26*18 | IGHV1-69*05 | 0.001 |
| IGHVF5S1-G26*19 | IGHV1-69*12 | 0.001 |
| IGHVF5S1-G26*20 | IGHV1-69*16 | 1e-04 |
| IGHVF5S1-G26*21 | IGHV1-69*11 | 1e-04 |
| IGHVF5S1-G26*22 | IGHV1-69*15 | 0.001 |
| IGHVF5S1-G27*01 | IGHV1-8*03 | 0.001 |
| IGHVF5S1-G27*02 | IGHV1-8*01 | 1e-04 |
| IGHVF5S1-G27*03 | IGHV1-8*02 | 0.001 |
| IGHVF5S1-G28*01 | IGHV1-46*04 | 0.001 |
| IGHVF5S1-G28*02 | IGHV1-46*03 | 0.001 |
| IGHVF5S1-G28*03 | IGHV1-46*01 | 0.001 |
| IGHVF5S1-G28*04 | IGHV1-46*02 | 1e-04 |
| IGHVF5S1-G29*01 | IGHV1-2*07 | 1e-04 |
| IGHVF5S1-G29*02 | IGHV1-2*04 | 1e-04 |
| IGHVF5S1-G29*03 | IGHV1-2*02 | 0.001 |
| IGHVF5S1-G29*04 | IGHV1-2*03 | 1e-04 |
| IGHVF5S1-G29*05 | IGHV1-2*01 | 1e-04 |
| IGHVF5S1-G29*06 | IGHV1-2*05 | 1e-04 |
| IGHVF5S1-G29*07 | IGHV1-2*06 | 0.001 |
| IGHVF5S1-G30*01 | IGHV1-18*04 | 0.001 |
| IGHVF5S1-G30*02 | IGHV1-18*01 | 0.001 |
| IGHVF5S1-G30*03 | IGHV1-18*03 | 1e-04 |
| IGHVF5S1-G31*01 | IGHV1-3*05 | 0.001 |
| IGHVF5S1-G31*02 | IGHV1-3*01 | 1e-05 |
| IGHVF5S1-G31*03 | IGHV1-3*04 | 0.001 |
| IGHVF5S1-G31*04 | IGHV1-3*02 | 1e-05 |
| IGHVF5S1-G31*05 | IGHV1-3*03 | 1e-04 |
| IGHVF6S1-G32*01 | IGHV2-26*04 | 1e-04 |
| IGHVF6S1-G32*02 | IGHV2-26*03 | 1e-04 |
| IGHVF6S1-G32*03 | IGHV2-26*01 | 1e-04 |
| IGHVF6S1-G32*04 | IGHV2-26*02 | 1e-04 |
| IGHVF6S1-G33*01 | IGHV2-5*08 | 0.001 |
| IGHVF6S1-G33*02 | IGHV2-5*01 | 0.001 |
| IGHVF6S1-G33*03 | IGHV2-5*02 | 5e-04 |
| IGHVF6S1-G33*04 | IGHV2-5*09 | 1e-04 |
| IGHVF6S1-G33*05 | IGHV2-5*05 | 0.001 |
| IGHVF6S1-G33*06 | IGHV2-5*06 | 1e-04 |

| | | |
|---|---|---|
| IGHVF6S1-G34*01 | IGHV2-70*12 | 1e-04 |
| IGHVF6S1-G34*02 | IGHV2-70*10 | 1e-04 |
| IGHVF6S1-G34*03 | IGHV2-70D*14 | 1e-04 |
| IGHVF6S1-G34*04 | IGHV2-70*16 | 1e-04 |
| IGHVF6S1-G34*05 | IGHV2-70*17 | 1e-04 |
| IGHVF6S1-G34*06 | IGHV2-70*04_A14G | 1e-04 |
| IGHVF6S1-G34*07 | IGHV2-70D*04/IGHV2-70*04 | 1e-04 |
| IGHVF6S1-G34*08 | IGHV2-70*01 | 1e-04 |
| IGHVF6S1-G34*09 | IGHV2-70*13 | 0.001 |
| IGHVF6S1-G34*10 | IGHV2-70*11 | 1e-04 |
| IGHVF6S1-G34*11 | IGHV2-70*15 | 1e-04 |
| IGHVF6S1-G34*12 | IGHV2-70*18 | 1e-04 |
| IGHVF6S1-G34*13 | IGHV2-70*19 | 1e-04 |
| IGHVF6S1-G34*14 | IGHV2-70*20 | 1e-04 |
| IGHVF7S1-G35*01 | IGHV6-1*02 | 5e-04 |
| IGHVF7S1-G35*02 | IGHV6-1*01 | 5e-04 |
| IGHVF7S1-G35*03 | IGHV6-1*01_T91C | 5e-04 |
| IGHVF8S1-G36*01 | IGHV4-4*08 | 1e-04 |
| IGHVF8S1-G36*02 | IGHV4-4*09 | 1e-04 |
| IGHVF8S1-G36*03 | IGHV4-59*08 | 1e-04 |
| IGHVF8S1-G36*04 | IGHV4-59*12 | 0.001 |
| IGHVF8S1-G36*05 | IGHV4-59*13 | 1e-04 |
| IGHVF8S1-G36*06 | IGHV4-59*11 | 0.001 |
| IGHVF8S1-G36*07 | IGHV4-59*07 | 1e-04 |
| IGHVF8S1-G36*08 | IGHV4-59*02 | 0.005 |
| IGHVF8S1-G36*09 | IGHV4-59*01 | 1e-04 |
| IGHVF8S1-G36*10 | IGHV4-59*01_G267A | 1e-04 |
| IGHVF8S1-G37*01 | IGHV4-59*10 | 1e-04 |
| IGHVF8S1-G37*02 | IGHV4-4*07 | 1e-04 |
| IGHVF8S1-G37*03 | IGHV4-4*07_A70G | 1e-04 |
| IGHVF8S1-G38*01 | IGHV4-34*09 | 1e-04 |
| IGHVF8S1-G38*02 | IGHV4-34*10 | 1e-04 |
| IGHVF8S1-G39*01 | IGHV4-34*11 | 1e-04 |
| IGHVF8S1-G39*02 | IGHV4-34*12 | 0.001 |
| IGHVF8S1-G39*03 | IGHV4-34*01 | 1e-04 |
| IGHVF8S1-G39*04 | IGHV4-34*02 | 0.002 |
| IGHVF8S1-G39*05 | IGHV4-34*04 | 1e-04 |
| IGHVF8S1-G39*06 | IGHV4-34*05 | 1e-04 |
| IGHVF8S1-G40*01 | IGHV4-4*01 | 1e-04 |
| IGHVF8S1-G40*02 | IGHV4-4*03 | 0.001 |
| IGHVF8S1-G40*03 | IGHV4-4*02 | 1e-04 |
| IGHVF8S1-G40*04 | IGHV4-4*10 | 1e-04 |
| IGHVF8S1-G41*01 | IGHV4-28*02 | 1e-05 |
| IGHVF8S1-G41*02 | IGHV4-28*06 | 1e-05 |
| IGHVF8S1-G41*03 | IGHV4-28*04 | 1e-05 |

| | | |
|---|---|---|
| IGHVF8S1-G41*04 | IGHV4-28*07 | 1e-05 |
| IGHVF8S1-G41*05 | IGHV4-28*05 | 1e-05 |
| IGHVF8S1-G41*06 | IGHV4-28*01 | 1e-05 |
| IGHVF8S1-G41*07 | IGHV4-28*03 | 1e-05 |
| IGHVF8S1-G42*01 | IGHV4-30-2*03 | 1e-04 |
| IGHVF8S1-G42*02 | IGHV4-30-4*07 | 1e-04 |
| IGHVF8S1-G42*03 | IGHV4-30-2*05 | 1e-04 |
| IGHVF8S1-G42*04 | IGHV4-30-2*06 | 5e-04 |
| IGHVF8S1-G42*05 | IGHV4-30-2*01_G70A | 1e-04 |
| IGHVF8S1-G42*06 | IGHV4-30-2*01 | 1e-04 |
| IGHVF8S1-G42*07 | IGHV4-30-2*01_C285T | 1e-04 |
| IGHVF8S1-G43*01 | IGHV4-30-4*02 | 1e-04 |
| IGHVF8S1-G43*02 | IGHV4-30-4*01_A70G_A107G | 1e-04 |
| IGHVF8S1-G43*03 | IGHV4-30-4*01 | 0.001 |
| IGHVF8S1-G43*04 | IGHV4-30-4*08 | 0.001 |
| IGHVF8S1-G44*01 | IGHV4-31*10 | 1e-04 |
| IGHVF8S1-G44*02 | IGHV4-31*11_G4C_G21C_C25T_A113C | 1e-04 |
| IGHVF8S1-G44*03 | IGHV4-31*11 | 1e-04 |
| IGHVF8S1-G44*04 | IGHV4-31*02 | 0.001 |
| IGHVF8S1-G44*05 | IGHV4-31*01 | 1e-04 |
| IGHVF8S1-G44*06 | IGHV4-31*03 | 1e-05 |
| IGHVF8S1-G45*01 | IGHV4-38-2*02_G246A | 0.001 |
| IGHVF8S1-G45*02 | IGHV4-38-2*01 | 1e-04 |
| IGHVF8S1-G45*03 | IGHV4-38-2*02 | 0.001 |
| IGHVF8S1-G45*04 | IGHV4-39*08 | 1e-04 |
| IGHVF8S1-G45*05 | IGHV4-39*02 | 1e-04 |
| IGHVF8S1-G45*06 | IGHV4-39*01_G315A | 0.005 |
| IGHVF8S1-G45*07 | IGHV4-39*01 | 0.001 |
| IGHVF8S1-G45*08 | IGHV4-39*01_A200C | 0.001 |
| IGHVF8S1-G45*09 | IGHV4-39*06 | 1e-04 |
| IGHVF8S1-G45*10 | IGHV4-39*07 | 0.001 |
| IGHVF8S1-G45*11 | IGHV4-39*09 | 1e-04 |
| IGHVF8S1-G46*01 | IGHV4-61*09 | 5e-04 |
| IGHVF8S1-G46*02 | IGHV4-61*02 | 0.001 |
| IGHVF8S1-G46*03 | IGHV4-61*11 | 5e-04 |
| IGHVF8S1-G46*04 | IGHV4-61*05 | 1e-04 |
| IGHVF8S1-G46*05 | IGHV4-61*10 | 1e-04 |
| IGHVF8S1-G46*06 | IGHV4-61*08 | 0.001 |
| IGHVF8S1-G46*07 | IGHV4-61*03 | 1e-04 |
| IGHVF8S1-G46*08 | IGHV4-61*01 | 1e-04 |
| IGHVF8S1-G46*09 | IGHV4-61*01_A41G | 1e-04 |

Table S1: **Allele cluster reference table.** The table contains the allele clusters for the full-length sequence germline set (S1) shown in Figure 2. The columns describe the alleles names according to the allele clusters naming scheme and the matching IUIS allele. The first column is the family, the second column is the allele cluster, the third column is the allele name, and the fourth column is the IUIS matching allele.
